# FLASH-RT does not affect chromosome translocations and junction structures beyond that of CONV-RT dose-rates

**DOI:** 10.1101/2023.03.27.534408

**Authors:** Paul G. Barghouth, Stavros Melemenidis, Pierre Montay-Gruel, Jonathan Ollivier, Vignesh Viswanathan, Patrik G. Jorge, Luis A. Soto, Brianna C. Lau, Cheyenne Sadeghi, Anushka Edlabadkar, Rakesh Manjappa, Jinghui Wang, Marie Le Bouteiller, Murat Surucu, Amy Yu, Karl Bush, Lawrie Skinner, Peter G. Maxim, Billy W. Loo, Charles L. Limoli, Marie-Catherine Vozenin, Richard L. Frock

## Abstract

The molecular and cellular mechanisms driving the enhanced therapeutic ratio of ultra-high dose-rate radiotherapy (FLASH-RT) over slower conventional (CONV-RT) radiotherapy dose-rate remain to be elucidated. However, attenuated DNA damage and transient oxygen depletion are among several proposed models. Here, we tested whether FLASH-RT under physioxic (4% O_2_) and hypoxic conditions (≤2% O_2_) reduces genome-wide translocations relative to CONV-RT and whether any differences identified revert under normoxic (21% O_2_) conditions. We employed high-throughput rejoin and genome-wide translocation sequencing (*HTGTS-JoinT-seq*), using *S. aureus* and *S. pyogenes* Cas9 “bait” DNA double strand breaks (DSBs), to measure differences in bait-proximal repair and their genome-wide translocations to “prey” DSBs generated by electron beam CONV-RT (0.08-0.13Gy/s) and FLASH-RT (1×10^2^-5×10^6^ Gy/s), under varying ionizing radiation (IR) doses and oxygen tensions. Normoxic and physioxic irradiation of HEK293T cells increased translocations at the cost of decreasing bait-proximal repair but were indistinguishable between CONV-RT and FLASH-RT. Although no apparent increase in chromosome translocations was observed with hypoxia-induced apoptosis, the combined decrease in oxygen tension with IR dose-rate modulation did not reveal significant differences in the level of translocations nor in their junction structures. Thus, Irrespective of oxygen tension, FLASH-RT produces translocations and junction structures at levels and proportions that are indistinguishable from CONV-RT.

---

FLASH radiotherapy (RT) is an ultrafast irradiation modality, generally more than two orders of magnitude faster than conventional dose rate (CONV) RT, demonstrating a significant enhancement of normal tissue sparing while still maintaining effective tumor control in nearly all cases [1-3]. To this point, physical beam parameters may influence the magnitude, or the actual benefit provided by ultra-high dose rate [4, 5]. Dose rate ≥40Gy/s is a putative threshold to achieve the “FLASH effect”, which has been observed mostly *in vivo* across different tissues of small and large animals (reviewed in: https://doi.org/10.1146/annurev-cancerbio-061421-022217, [3]). Early human clinical trials have also demonstrated feasibility of delivering FLASH dose rates [3, 6]. Despite the near consensus on the benefit of FLASH-RT over CONV-RT dose rates, the mechanism explaining the FLASH effect remains to be elucidated.

Multiple hypotheses have been proposed to explain the FLASH effect which include lipid peroxidation, mitochondrial metabolic differences, and metabolic hibernation as potential driving factors (reviewed in: https://doi.org/10.1146/annurev-cancerbio-061421-022217, [3]). Other hypotheses include a near instantaneous radioprotective hypoxic state afforded by FLASH-RT [7, 8] to ultimately suppress DNA damage. In this context, DNA double strand break (DSB) measurements have largely been limited to measuring the kinetics of DNA damage response intermediates in cells and tissues [9, 10] or by measuring DNA strand breaks from supercoiled plasmids [11]. Therefore, to determine whether the FLASH effect imparts any changes to DSB repair or pathway choice, a rigorous cellular genome-wide translocation assay that can measure DSBs from ionizing radiation (IR) is necessary.

High-Throughput Genome-wide Translocation Sequencing (*HTGTS*) [12] and its Linear Amplification Mediated (LAM) protocol [13, 14], leverages the generation of a fixed “bait” DSB to capture “prey” DSBs via their chromosome translocation to each other and maps the resulting prey end of the junction (or joint) at single nucleotide resolution. *LAM-HTGTS* has been used extensively to quantify translocations generated from designer endonucleases [13, 15-17], physiologic DSBs [18-21] and IR [13]. ReJoin and Translocation sequencing (*JoinT-seq*) built upon the *LAM-HTGTS* platform by additionally quantifying proximal repair outcomes (e.g. deletions) of the bait “Breaksite” along with genome-wide translocations and recently described DSB repair in the context of core nonhomologous end joining (NHEJ) deficiency of G1-arrested cells [21].

Here, *HTGTS-JoinT-seq* is employed to identify alterations in translocations and/or proximal repair affected by irradiation dose-rate under normoxic, physioxic, and hypoxic oxygen tensions in 293T cells. Although enrichment of translocations is IR dose-dependent and is separately attenuated by hypoxic oxygen tensions, FLASH-RT does not confer any significant decrease in translocations or alteration of junction structures beyond that imposed by CONV-RT.

## Materials and Methods

### Cell culture and Cas9 plasmids

HEK293T cells were cultured at 37°C, 5% CO_2_ using DMEM supplemented with glutamine, 10% FCS and 0.5% penicillin/streptomycin. The following *S. aureus* Cas9 plasmids were kindly provided by Manuel Gonçalves (Leiden): BA15_pCAG.SaCas9.rBGpA (SaCas9 nuclease) and AV85_pSa-gRAG1.1 (encoding RAG1-specific Sa-gRNA1.1) [15, 17]. *S pyogenes* Cas9 plasmid, pX330-U6-Chimeric_BB-CBh-hSpCas9, was obtained from Addgene (#42230). RAG1B gRNA cloning and pCMX-eGFP have been previously described [13]. See supplement for more details.

### Oxygen tension

SaCas9:RAG1.1-transfected cells were either kept at normoxia (21% O2) or placed into a controlled atmosphere chamber (InVivo2 Hypoxic Workstation at Stanford and Biospherix hypoxia hood at CHUV) at hypoxic oxygen tensions of 4%, 2%, or 0.5% O_2_ set to 37°C. A portable hypoxic chamber OxyGenie (Baker) was used to transport cells to and from the radiation facility at Stanford. See supplement for more details.

### Irradiation Exposure

SaCas9:RAG1.1 experiments were performed at Stanford University using a Varian Trilogy medical LINAC (Varian Medical Systems, Inc., Palo Alto, CA) configured for ultra-high dose rate electron beam delivery. SpCas9:RAG1B experiments and clonogenic survival were performed at CHUV Lausanne University Hospital using Oriatron eRT6 electron beam LINAC (PMB Alcen). See supplement for beam parameters.

### Flow cytometry analysis

Cells were measured for GFP (transfection) and 7AAD (BD) (cell death) at end points for *HTGTS-JoinT-seq* or for PI/FITC-Annexin V (Biolegend) staining for apoptosis detection. In all cases, untransfected cells were subjected to no, single, and double staining to optimize gating.

### HTGTS-JoinT-seq

12μg input DNA complexity was used to enrich for junction libraries; primers for SaCas9:RAG1.1 [15, 17] and SpCas9:RAG1B [13, 16] bait DSBs were previously described. SpCas9:RAG1B libraries were sequenced using MiSeq (250PE) and normalized to 214,700 sequence reads. SaCas9:RAG1.1 libraries were sequenced by NovaSeq (150PE) and normalized to 904,846 sequence reads. Sequence data were processed as described [21] and aligned to hg38 genome build. MACS2 identified hotspots as described [13].

## Results

### CONV-RT increases genome-wide translocations

We first assayed proximal and genome-wide DSB repair outcomes influenced only by CONV-RT. For all experiments described, we distinguished translocations versus bait-proximal repair (termed as the “Breaksite”) by separating recovered prey junctions aligned outside versus inside, respectively, a 500kb window flanking each side of the bait DSB since spreading of the phosphorylated H2AX damage signal covers a range of hundreds of kilobases up to several megabases [22-24]. We employed the *S. aureus* Cas9 bait DSB targeting the RAG1 locus on chromosome 11 using the RAG1.1 guide RNA (SaCas9:RAG1.1) in 293T cells [15, 17], which has a higher substrate turnover rate in comparison to the commonly used *S. pyogenes* Cas9 [25], and assayed for repair outcome differences over a 24 hour period after 10 Gy exposure from a CONV-RT electron beam clinical linear accelerator (LINAC; Stanford) [26, 27] (**Fig. 1A**). Cell death was minimally increased with irradiation, and the cell fractions with transfected GFP reporter/Cas9 plasmids were relatively high by flow cytometry (**Fig. 1B**) prior to collection for *JoinT-seq* library preparation and sequencing.

**Fig. 1:**
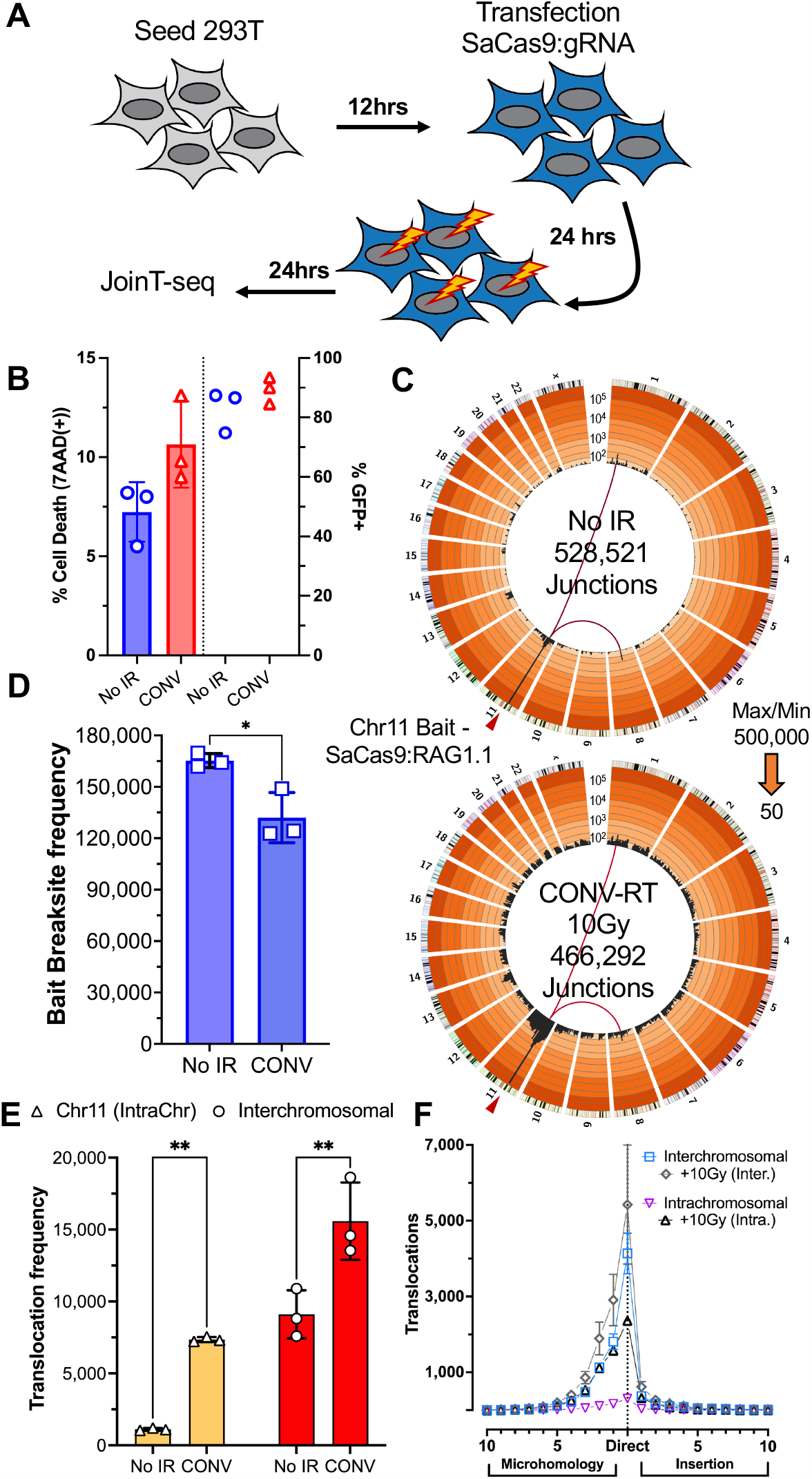
Intrachromosomal translocations are dramatically increased after CONV-RT. (A) Strategy for SaCas9:gRNA delivery, electron beam irradiation and collection for *JoinT-seq* analysis. (B) Cell death and transfection efficiency measures by flow cytometry. (C) Circos plots describing translocations in 5Mb binned regions across each chromosome (black bars) displayed on a custom 1,2,5 iteration log scale with increasingly darker orange coloring indicating greater orders of magnitude scaling; Max/Min ranges are indicated. Red triangle indicates position of the bait DSB bin and inclusive of the bait Breaksite window (N=3 each). (D) Junction frequencies within the bait Breaksite; unpaired t-test: *P < 0.05. (E) Intra- and interchromosomal translocation frequencies; two-way ANOVA with Šidák posttest: **P < 0.01. (F) Translocation junction structure distributions.

Approximately half a million total junctions were recovered for the non-irradiated (No IR) control (**Table S1**); two previously identified RAG1.1 off-target hotspots on chromosomes 1 and 8 [15, 17] were identified again (**Fig. 1C**). Deeper analysis revealed ∼94% of junctions aligned within the bait Breaksite (**Fig. 1C, D; Table S1**), with most of the total (∼90%) generating small (≤25bp) deletions as a direct result of rejoining bait DSB ends (**Fig. S1A, B**) [21]. Beyond the Breaksite window, intrachromosomal translocations were ∼150-fold less frequent; however, interchromosomal translocations were ∼8-fold greater than their intrachromosomal counterpart, averaging ∼9,100 junctions (**Fig. 1E; Table S1**). Both translocation subgroups predominantly harbored direct, with some short microhomology (MH), joints, consistent with NHEJ utilization (**Fig. 1F**) [28]; although, translocations significantly relied more on long MHs in contrast to small deletions (∼3% vs. ∼13%)(**Fig. S1C**).

CONV-RT resulted in a total net loss in recovered junctions by 12% and specifically affected bait Breaksite junctions (**Fig. 1C, D; Fig. S1B; Table S1**). Correspondingly, CONV-RT increased genome-wide translocations 2.1-fold, totaling ∼69,000 translocations (∼22,900 on average) (**Fig. 1C, E; Table S1**). Crucially, intrachromosomal translocations outgained interchromosomal translocations by nearly 4-fold, rendering the chromosome harboring the bait DSB a hotspot for translocation (**Fig. 1C, E; Table S1**). Despite the increased genome-wide translocations, CONV-RT did not alter their junction structure proportions (**Fig. S1D**), suggesting the bulk of translocations from both spontaneous DSBs and IR-generated DSBs are repaired in a similar manner. Thus, we conclude CONV-RT from electron beam LINAC (**Table S2**) imparts a similar translocation enrichment pattern as previously observed using X-Ray sources [13, 29].

### Translocations are not increased under chronic hypoxia

We next sought to describe genome-wide translocations in the context of varied oxygen tensions. Culture of 293T cells at physiologic (2% O_2_) and pathologic (0.5% O_2_) hypoxic oxygen tensions over 24 hours did not reveal a dramatic change in cell death (<10%) relative to the normoxic (21% O_2_; air) environment. However, cell death was significantly increased over a 48-hour period, particularly for 0.5% O_2_ oxygen tensions, affecting 45-75% of the population (**Fig. S2A**), suggesting potentially increased levels of translocations. SaCas9:Rag1.1 was transiently delivered into 293T cells and split into three oxygen tensions–21% O_2_, 2% O_2_, and 0.5% O_2_—and assayed for repair after 48 hours (**Fig. 2A**). Transfection was efficient in all three conditions despite reproducing an O_2_ level-dependent effect on cell death (**Fig. S2B**). Total recovered junctions for normoxic cells were nearly 600,000 but were progressively decreased by 15-25% as oxygen tension attenuated (**Fig. 2B; Table S1**). In striking contrast to CONV-RT (see Fig. 1), junctions from both translocations and the bait Breaksite were decreased 10-25% under hypoxic conditions (**Fig. 2C, D; Table S1**). Hypoxia did not affect translocation junction structure distributions (**Fig. S2C**) but small deletions in the Breaksite used progressively less (3-5%) polymerase-mediated insertions (**Fig. S2D**). Despite the high level of cell death from chronic hypoxia, we could not identify any discernible increase in chromosome translocations. We revisited the type of cell death occurring at 48 hours of hypoxia and found most of the dying cells were apoptotic (**Fig. S2E, F**). To confirm whether cycling cells were specifically impacted by hypoxia as concluded from earlier studies [30], we measured for apoptosis in cycling and G1-arrested Abelson kinase-transformed progenitor B cells [21] with decreasing O_2_ levels over 48 hours and found cycling cells, but not the G1-arrested population, were subjected to increasing apoptosis as a function of decreasing O_2_ (**Fig. S2F**). Thus, we conclude chronic pathologic hypoxia induces apoptosis in cycling cells but curiously does not increase the level of translocations.

**Fig. 2:**
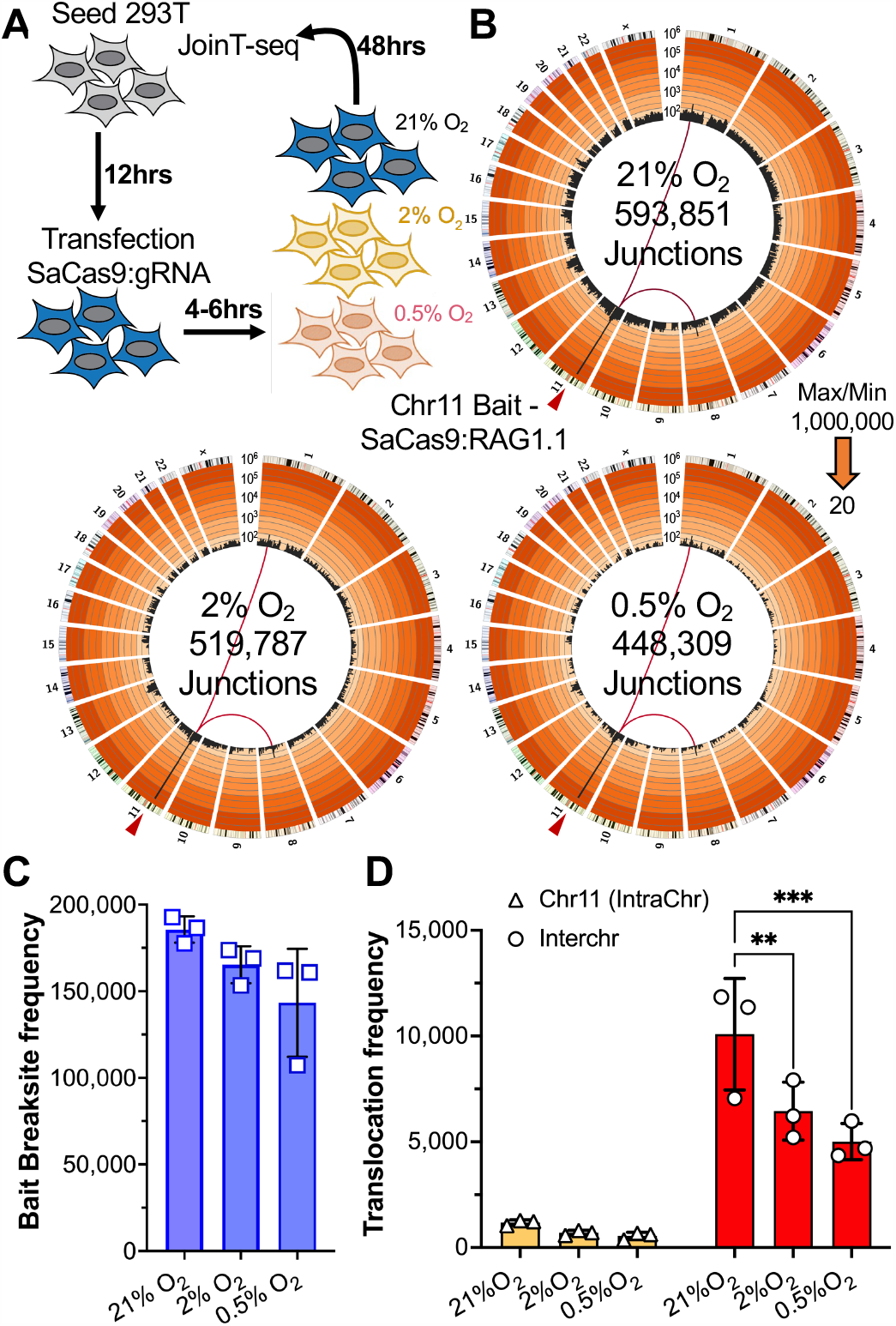
Chronic hypoxia does not induce translocations. (A) Strategy to detect translocations under different oxygen tensions. (B) Circos plots as described in Figure 1 (N=3 each). Max/Min range is indicated. (C) Bait Breaksite frequencies; one-way ANOVA with Dunnett’s posttest: no significance below P < 0.05. (D) Translocation frequencies; two-way ANOVA with Dunnett’s posttest: **P < 0.01, ***P < 0.001.

### Translocations are undistinguished between CONV-RT and FLASH-RT

To discern FLASH dose-rate effects on DSB repair, we first assayed repair at physioxic oxygen tension (4% O_2_) employing the more commonly used *S. pyogenes* Cas9 (SpCas9), and gRNA bait, RAG1B [13] targeting a DSB 45bp away from the SaCas9:RAG1.1 DSB site. Here, transfected 293T cells (**Fig. S3A**) were transitioned to 4% O_2_ environment and irradiated with 10 Gy using either an X-Ray tube (0.06Gy/s) or an eRT6 electron beam LINAC set for CONV-RT (0.21Gy/s) or FLASH-RT (1,000Gy/s) (CHUV) (**Fig. 3A**). Recovered junctions for the control totaled just over 70,000 and RAG1B off-target DSB hotspots were again found on chromosomes 14 and 4 (**Fig. 3A; Table S1**), as previously reported [13, 16]. Analogous to the SaCas9:RAG1.1 bait, the SpCas9:RAG1B bait Breaksite consisted of ∼90% of total junctions followed by interchromosomal and intrachromosomal translocations (**Fig. 3A-C; Table S1**). Similarly, all irradiated samples resulted in fewer junctions recovered (25-33% less than control), with a 2-fold decrease in bait Breaksite repair and a characteristic increase of intrachromosomal translocations that, combined with interchromosomal translocation gains, led to a ∼6-fold increase in total translocations (**Fig. 3A-C; Table S1**). Crucially, we did not observe any significant differences in translocation numbers or junction structures between radiation sources or dose-rate modalities at 4% O_2_ (**Fig. 3C-D; Table S1**), and clonogenic survival did not reveal any significant difference between CONV-RT and FLASH-RT (**Fig. S3B; Table S3**). We conclude IR dose rate modulation does not impact translocations and cell survival for 293T cells at physioxic oxygen tension.

**Fig. 3:**
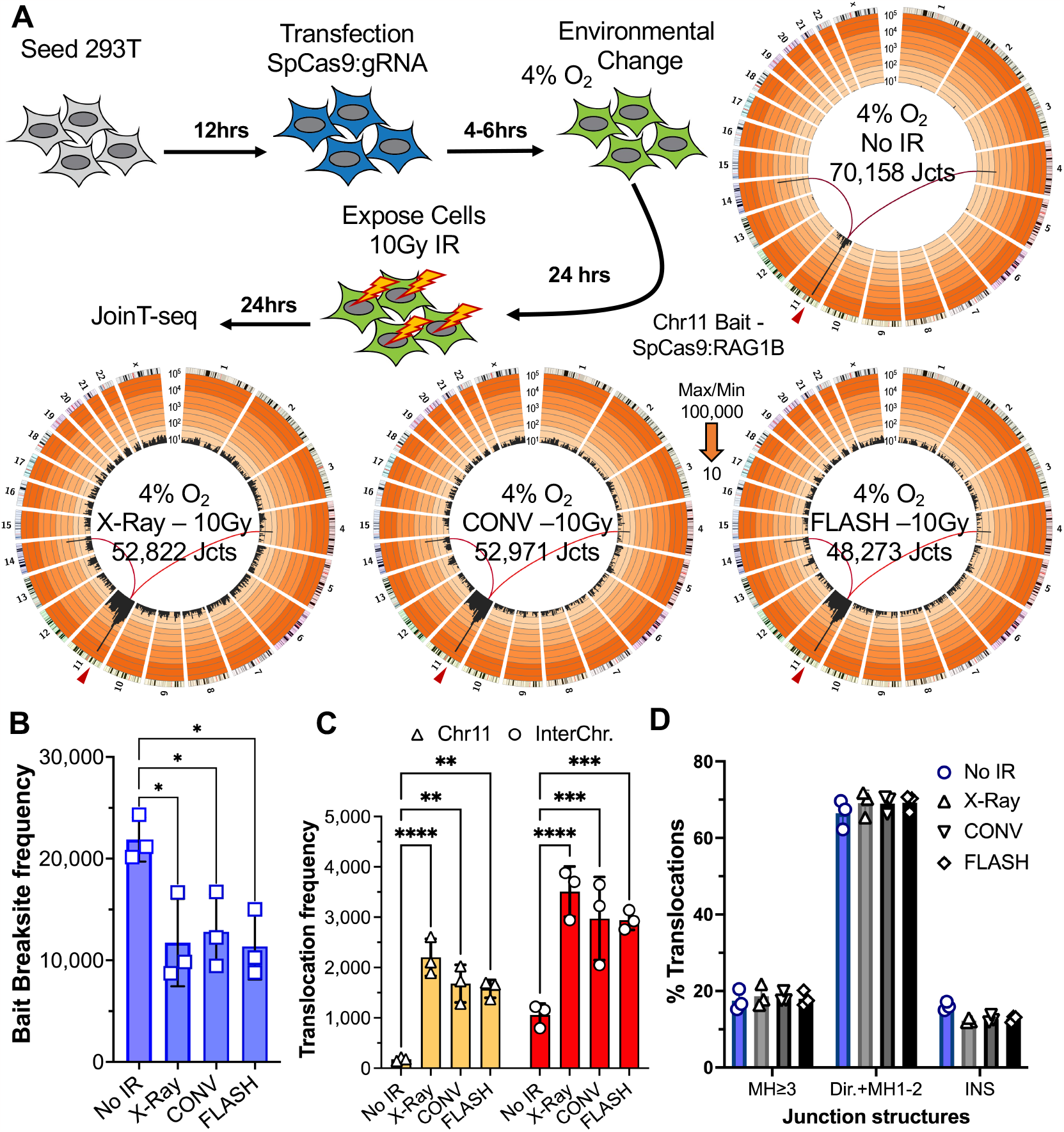
Translocations remain undifferentiated between dose rates at 4% O_2_. (A) Physioxic irradiation strategy and resulting circos plots from control, X-Ray, CONV, and FLASH irradiated samples (N=3 each). (B) Bait-proximal junction frequencies; One-way ANOVA with Dunnett’s posttest: *P < 0.05. (C) Translocation frequencies; **P < 0.01, ***P < 0.001, ****P < 0.0001. (D) Translocation junction structure distributions split into three groups: long microhomologies (MH≥3), direct with short MHs (Dir+MH1-2), and insertions (INS); two-way ANOVA with Tukey posttest: no significance below P < 0.05.

Finally, we sought to identify any potential translocation differences between CONV-RT and FLASH-RT at pathologic hypoxic oxygen tension (0.5% O_2_), where the FLASH effect may be most pronounced. SaCas9:RAG1.1 transfected 293T cells cultured in normoxic or hypoxic conditions, transported either in tubes or in a portable environment-controlled apparatus (i.e., OxyGenie), and irradiated under CONV-RT or FLASH-RT settings (Stanford) (**Figs. 4A; S4A-C**)(see methods and supplement). Gafchromic film dosimetry confirmed CONV and FLASH dose-rates delivered per replication (**Fig. 4B; Table S2**). For all conditions assayed (oxygen tension and dose-rate; 4 irradiation doses and 1 non-irradiated control for each combination), transfection efficiency was consistently high and cell death for hypoxic cultures ranged 30-50% (**Fig. S4D**). Total recovered junctions across normoxic controls ranged between ∼452,000-677,000 with hypoxia decreasing recovered junctions by 5-12% (**Figs. S5A, S6A; Table S1**); overall, irradiation depressed total recovered junctions (**Figs. S5A-C**; **S6A-C; Table S1**). As predicted, the fraction of translocations was progressively increased as a function of IR dose (**Figs. 4C, D; S5A-C; S6A-C; Table S1**). In this regard, irradiation under pathologic hypoxia also yielded increased translocations, though the rate increase with absorbed dose increase was shallower (**Fig. 4C, D; Table S1**). Hypoxia together with irradiation did not alter translocation junction structure distributions (**Fig. S4E**). Importantly, translocation numbers and junction structures between CONV-RT and FLASH-RT at comparable absorbed doses and oxygen tensions were indistinguishable (**Fig. 4C, D; S3E; Table S1**). We conclude that irradiation dose rate at pathologic hypoxic oxygen tensions does not affect translocation formation and frequency.

**Fig. 4.**
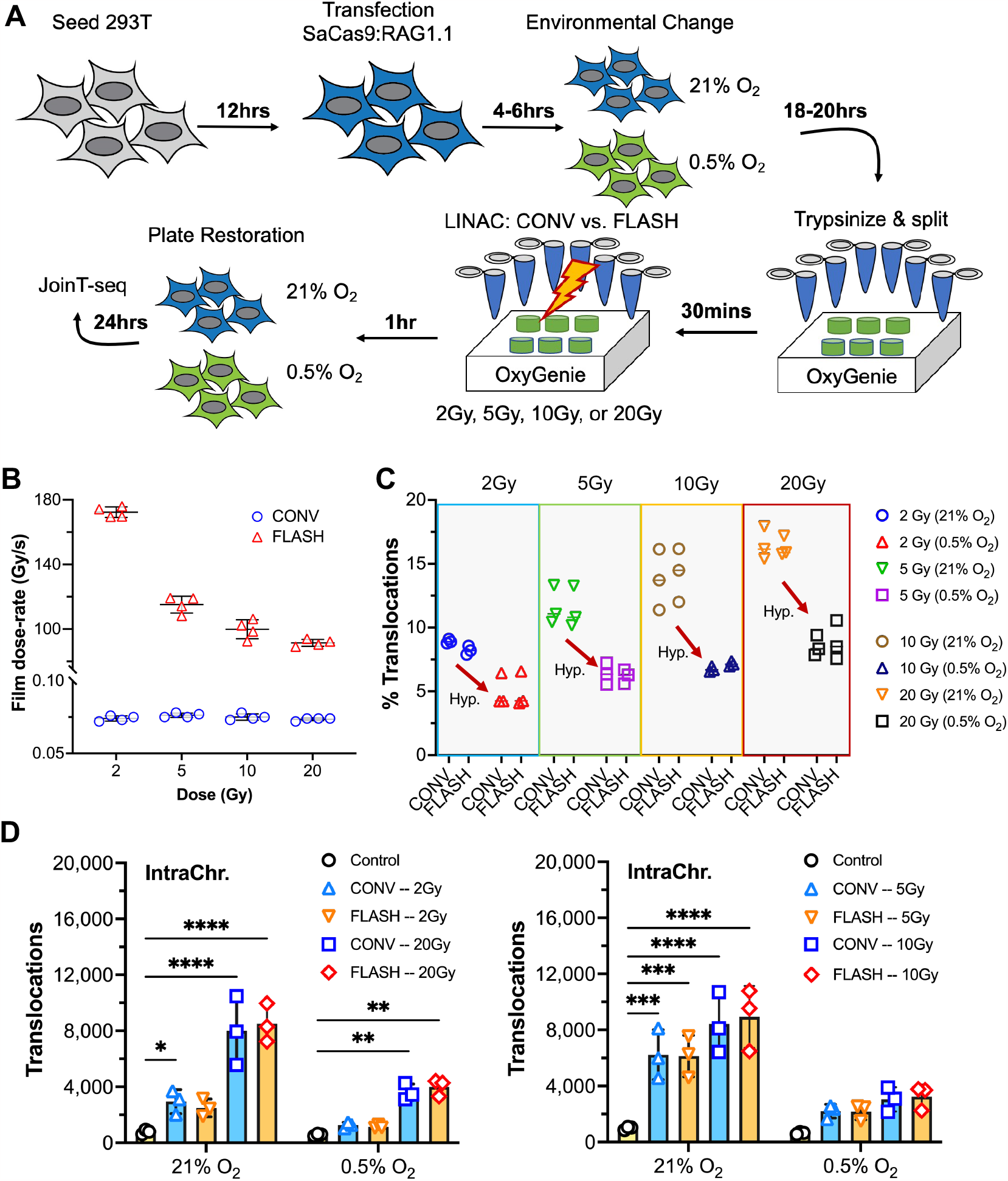
FLASH-RT and CONV-RT display similar translocation increases with increasing dose. (A) Strategy for normoxic and hypoxic irradiation measures for JoinT-seq. Cells in both environments were trypsinized and maintained in their respective oxygen tensions during irradiation. (B) Film dose rates plotted for each experiment (N=3) (C) Percent of translocations for each dose rate/environment/absorbed dose over a 2 Gy–20 Gy range. (D) Intrachromosomal translocation frequency gains with increasing absorbed IR dose for the 2/20 Gy and the 5/10Gy experimental data sets; two-way ANOVA with Dunnett’s posttest: *P < 0.05, **P < 0.01, ***P < 0.001, ****P < 0.0001.

## Discussion

The goal of this study was to determine whether any molecular differences in DSB repair or translocations could be identified between ionizing radiation dose-rate extremes and if those differences could help reveal mechanisms leading to the “FLASH effect”. The premise was that potential molecular alterations in DSB repair outcomes afforded by FLASH dose rates at low oxygen tensions may not significantly impact cell viability, given that the 293T line is SV40 large T antigen transformed, but that alterations to the formation of DSBs and translocation would be discerned irrespective of normal or transformed status. Using *HTGTS-JoinT-seq* [21], we found IR dose-rate to have no significant impact on translocations and proximal repair beyond the intended absorbed dose, across a wide range of oxygen tensions, and as determined from two different FLASH irradiators. While these data do not formally rule out potential dose-rate dependent differences in the yields of other radiolytic lesions (single strand breaks, base damage), present results clearly indicate that the induction and repair of DSB lesions is not dose-rate dependent with regard to the cells and range of dose rate assayed here. Furthermore, based on these and other available data (https://doi.org/10.1146/annurev-cancerbio-061421-022217), the potential role of DNA damage and repair as an underlying mechanism of the FLASH effect seems implausible.

We also discovered oxygen tensions down to 0.5% O_2_ did not induce substantial apoptosis in 293T cells until after 24 hours of low oxygen tensions. Here, the replicating cell cycle phases (S-G2/M) are implicated in the cell death phenotype since G1-arrest did not significantly increase apoptosis detection. The cell death mechanism and response observed under chronic hypoxia is similar to what has been previously described in 24-hour cultures, but under extremely low oxygen tension (<0.1% O_2_), resulting initially in a stressed, but sustained, replication from a limiting nucleotide pool with eventual stalling and, ultimately, replisome unloading beyond 12 hours of extreme hypoxia [30-32]. In this context, hypoxia-mediated activation of the DNA damage kinases and phosphorylation of H2AX (γ-H2AX) [30, 33, 34] are necessary to stabilize stalled replication forks and to suppress apoptosis. Yet, despite a high level of apoptosis observed at 48 hours of low oxygen tensions in our studies, additional translocations were not detected.

The above findings agree with early-stage apoptosis observations demonstrating pan-nuclear or nuclear-peripheral γ-H2AX staining that are not enriched with DSB repair markers, unlike with individual DNA damage foci [35, 36] or micronuclei [37, 38]. However, further insight comes from a recent study that determined apoptotic DNA fragmentation generates extrachromosomal circular DNA elements (eccDNAs) with singular DNA alignments and size distributions varying in units of nucleosome occupancy [39]. In this regard, genome-wide circularization of DNA fragments, resulting in the formation of potent immunostimulants for dendritic cells and macrophages, requires DNA ligase III, but not the NHEJ ligase, DNA ligase IV [39]. Therefore, hypoxia-induced apoptotic eccDNA precursors would not be translocation substrates for Cas9 bait DSBs in human cells [28]. We speculate repair pathway incompatibility along with other contributing factors—(1) frequency and proximity of eccDNA precursor ends to each other relative to the bait DSB, (2) end accessibility differences, and (3) caspase-mediated cleavage of repair proteins [13, 15, 21, 40, 41]—are at play to promote the cell death program and to stimulate phagocytosis of apoptotic bodies. A deeper investigation into apoptotic DNA recombination will be necessary to better understand mechanisms driving eccDNA biogenesis and any potential role eccDNAs have in explaining the FLASH effect *in vivo*.

## Author Contributions

PGB, CLL, M-CV, RLF designed the study. PGB, SM, PM-G, JO, VV, PGJ performed experiments. PGB, CS, AE, MLB, RLF analyzed *HTGTS-JoinT-seq* data. SM, LAS, BCL performed and SM, RM, JW optimized Stanford dosimetry. PGJ, MS, AY, KB, LS, PGM, BWL Jr., CLL, M-CV supervised irradiations, machine, and dosimetry Q/A. RLF supervised the research. PGB and RLF wrote the manuscript. BWL Jr., CLL, and M-CV commented on and edited the manuscript.

## Data Availability

Illumina sequencing data are available through GEO (GSE227466).

## Acknowledgements

The authors thank Manuel Gonçalves (LUMC) for providing the SaCas9:Rag1.1 vectors, Erinn Rankin lab and Baker Ruskinn for the OxyGenie device, and members of the RLF, BWL, CLL, and M-CV labs for discussion and feedback. This work was supported by the NIH Ruth L. Kirschstein National Research Service Award fellowship T32CA121940 (to PGB), the Radiation Research Foundation Career Development Award (to RLF) and by the NCI grant P01CA244091 (to BWL Jr., M-CV and CLL).

