## Supplement for "FLASH-RT does not affect chromosome translocations and junction structures beyond that of CONV-RT dose-rates"

**Supplemental methods**

**Cell transfection.** HEK293T cells were transfected using 1 mg/mL 25 kDa linear polycation polyethyleneimine (PEI) solution at pH of 7.4. 293T cells were seeded at ~1.3 x 10^6 cells/mL into a 10cm dish containing 9mL of fresh DMEM and incubated for 12 hours prior to transfection. For SaCas9:RAG1.1 transfection, a ratio of 1:3 DNA to PEI was generated; specifically, 11μg DNA to 33μL PEI. An equimolar plasmid mixture containing 74% SaCas9 nuclease, 16% Sa-RAG1.1 gRNA, and 10% eGFP plasmids were diluted in a 150mM NaCl to a final volume of 500μL and vortexed for 10 seconds. In a separate tube, 33μL PEI was diluted into 150mM NaCl to a final volume of 500μL and vortexed. Importantly, the transfection mixture was made by adding 500μL of PEI-NaCl mixture into the tube containing 500μL DNA-NaCl and immediately vortexed for 30 seconds. Thereafter, the transfection mixture was incubated for 30 minutes to allow for PEI-DNA complexes to form. The resulting 1mL of DNA:PEI mixture was added, drop-wise, into the 9mL of culture media of the pre-seeded 10mm dish and was then gently rocked to mix the transfection reagent homogeneously within the culture media. FuGENE (Promega) transfections were performed for SpCas9:RAG1B experiments in HEK293T cells according to the manufacturer's protocol. Briefly, 20μg of total plasmid (10% pCMX-eGFP) was transfected with FuGENE in a 2:1 FuGENE:DNA ratio.

**Culture of *v-Abl* cells:** The Abelson kinase-transformed (*v-Abl*) murine progenitor B cell line “WT A” clone [21] was cultured at 37°C and 5% CO_2_ in RPMI supplemented with 10% FCS, 50 U/mL penicillin/streptomycin (Gemini), 2 mM L-glutamine (Gemini), 1× MEM-NEAA (Lonza), 1 mM sodium pyruvate (Cytiva), 50 μM 2-mercaptoethanol (Sigma), and 20 mM Hepes (pH 7.4)(Cytiva). G1/G0-arrest was induced using 3μM STI-571 (TCI), placed into 0.5% O_2_ after 48 hours (when fully arrested) and assayed for viability after another 48 hours. Asynchronous *v-Abl* cells were seeded in fresh medium and cultured for 48 hours at 0.5% O_2_.

**Clonogenic survival.** 293T cells were cultured in monolayer with complete medium containing DMEM (Dulbecco’s Modified Eagle Medium) + GlutaMAX (4.5g/L D-Glucose, Pyruvate) (31966-021, Gibco) and supplemented with 10% FBS for cell culture (F7524, Sigma). Cells were incubated in a hypoxia chamber (Biospherix) at 4% O_2_, 5% CO_2_ 24 hours before the irradiation. The next day, in the hypoxia chamber, cells were harvested, counted, and transferred in 2 mL Eppendorf tubes for irradiation. Tubes (500,000 cells in 1 mL medium) were placed in a water tank before irradiation at 2, 4, 6, and 8 Gy using FLASH and CONV settings reported in **Table S3**. Next, cells were plated in triplicate at a concentration of 200 – 4000 cells/well in a 6-well cell culture plate and incubated at 37°C, 21% O_2_ & 5% CO_2_ until colonies were visible. 2 weeks after irradiation, colonies were fixed, stained with crystal-violet 0.1% (Sigma) and counted. Plating efficiency and percentage survival was determined. On GraphPad Prism, the linear quadratic model was used to fit survival curves.

**Oxygen tension control for *HTGTS-JoinT-seq*.** Hypoxia treatment time frame was dependent on experimental requirements, either 24 or 48 hours. *HTGTS-JoinT-seq* experiments were terminated by the 48-54th hour post transfection, assessed for transfection efficiency via flow cytometry analysis (eGFP+ >50%) and genomic DNA isolated from cells. SpCas9:RAG1B-transfected cells were kept at physioxic tension (4% O_2_) in a different controlled atmospheric chamber (Biospherix C-Chamber with Biospherix ProOx C21 controller at Lausanne) for 48 hours prior to sample collection and genomic DNA isolation except for irradiation at the +24-hour controlled oxygen tension period (<1 hour duration).

**Transport and handling of normoxic, physioxic, and hypoxic cells to irradiators.** We set up two cohort replication sets for SaCas9:RAG1.1 irradiation experiments, 2 Gy/20 Gy and 5 Gy/10Gy (CONV-RT vs FLASH-RT), cultured either in 21% O_2_ or 0.5% O_2_, which included mock irradiated control for each condition. To maintain oxygen levels at 0.5% O_2_ during irradiation +24 hours ∆O_2_, the portable hypoxic chamber OxyGenie (Baker) was used to transport cells to and from the radiation facility. OxyGenie provides a constant input of gas from its miniature gas cylinder filling station to the cell that resides in airtight OxyGenie silicone cell culture well (~30mm well). When assembled, the culture cassettes hold an airtight seal for greater ~15 minutes when disconnected from the system [42].

Trypsinization (trypsin and 1xPBS) reagents were equilibrated in the hypoxia chamber (0.5% O_2_) for 5 hours prior to their use. Cells were trypsinized for 5 minutes, neutralized in 2mL 1xPBS, and stored on ice at their respective oxygen tensions before spun down at 1500x g for 5 minutes and resuspended with the original DMEM plus transfection reagent. Prior to placing normoxic cells into 1.5mL Eppendorf tubes and hypoxic cells into the OxyGenie cassettes, resuspended cells were pooled together to homogenize transfected cells and then distributed at 1mL per tube or OxyGenie cassette. Hypoxic cells were removed from the chamber and connected to the OxyGenie airflow system. To minimize cell stress from extended suspension, cells were kept at 4^o^C. Post irradiation, cells were resuspended, and volume was restored in a 10cm dish containing ~9mL DMEM plus transfection reagent. Cells were returned to their respective incubators for an additional 24 hours prior to collection at +48-56 hours, flow cytometry analysis, and genomic DNA isolation. A similar trypsinization and replating method was used for SpCas9:RAG1B physioxic experiments, but with transport of 4% O_2_ cells controlled in pre-equilibrated solutions and transport/irradiation in tightly sealed tubes.

**Irradiation Exposure.** SaCas9:RAG1.1 transfected cells were irradiated in the Stanford University Department of Radiation Oncology at CONV-RT (0.08Gy/s) or FLASH-RT (100-180Gy/s) 24 hours post oxygen tension equilibration, using a Varian Trilogy medical LINAC (Varian Medical Systems, Inc., Palo Alto, CA). Clinical mode was used for CONV-RT but configuration for FLASH-RT was as described elsewhere [10, 27, 43, 44]. In brief, a decommissioned electron energy board was tuned to achieve FLASH beam parameters with samples placed inside the treatment head (closer to source/scattering foil). For CONV-RT a clinical electron beam was used (16 MeV) and the samples were placed farther away from the source. The mean energies of the electron beams were calculated from the 50% of the maximum (R50) and were 15.7 MeV for CONV-RT and 16.6 MeV for FLASH-RT [45]. The delivery of the dose was confirmed by Gafchromic EBT3 films (Ashland, USA) for each sample. All film dosimetry was performed in accordance with the task group report 235 of the American Association of Physicists in Medicine [46]. SpCas9:RAG1B transfected cells were irradiated at Lausanne University Hospital using an X-Ray tube (0.06Gy/s) or a prototype Oriatron eRT6 5.5 MeV electron beam LINAC (PMB Alcen) for CONV (0.21Gy/s) or FLASH (1,000Gy/s). See **Tables S2, S3** for CONV-RT and FLASH-RT beam parameter details for each irradiator.

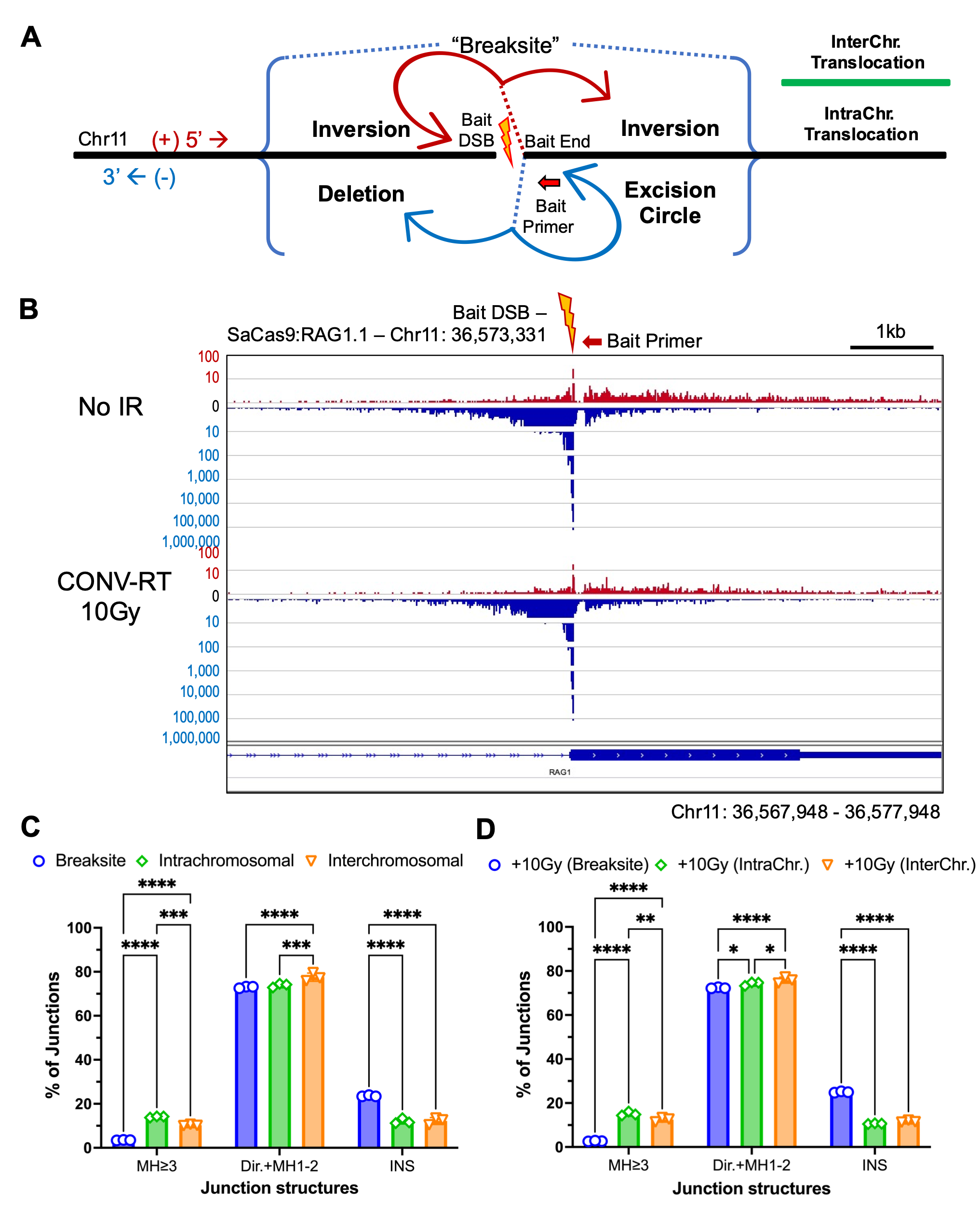

**Fig. S1: SaCas9:RAG1.1 Bait Breaksite.** (A) Proximal repair outcomes as previously described (Frock et al., 2015). Note the red bait primer is in the minus (blue) orientation. (B) IGV plot depicting massive enrichment of small deletions. (C) Junction structure distributions between the Breaksite, intra-, and interchromosomal translocations. Long microhomologies (MH-≥3), Direct/short MHs (Dir+MH1-2) and insertions (INS) are split across each subgroup. (D) Junction structures of the same subgroups as in (C) but with 10 Gy CONV-IR. Two-way ANOVA with Šidák posttest: *P < 0.05, **P < 0.01, ***P < 0.001, ****P < 0.0001.

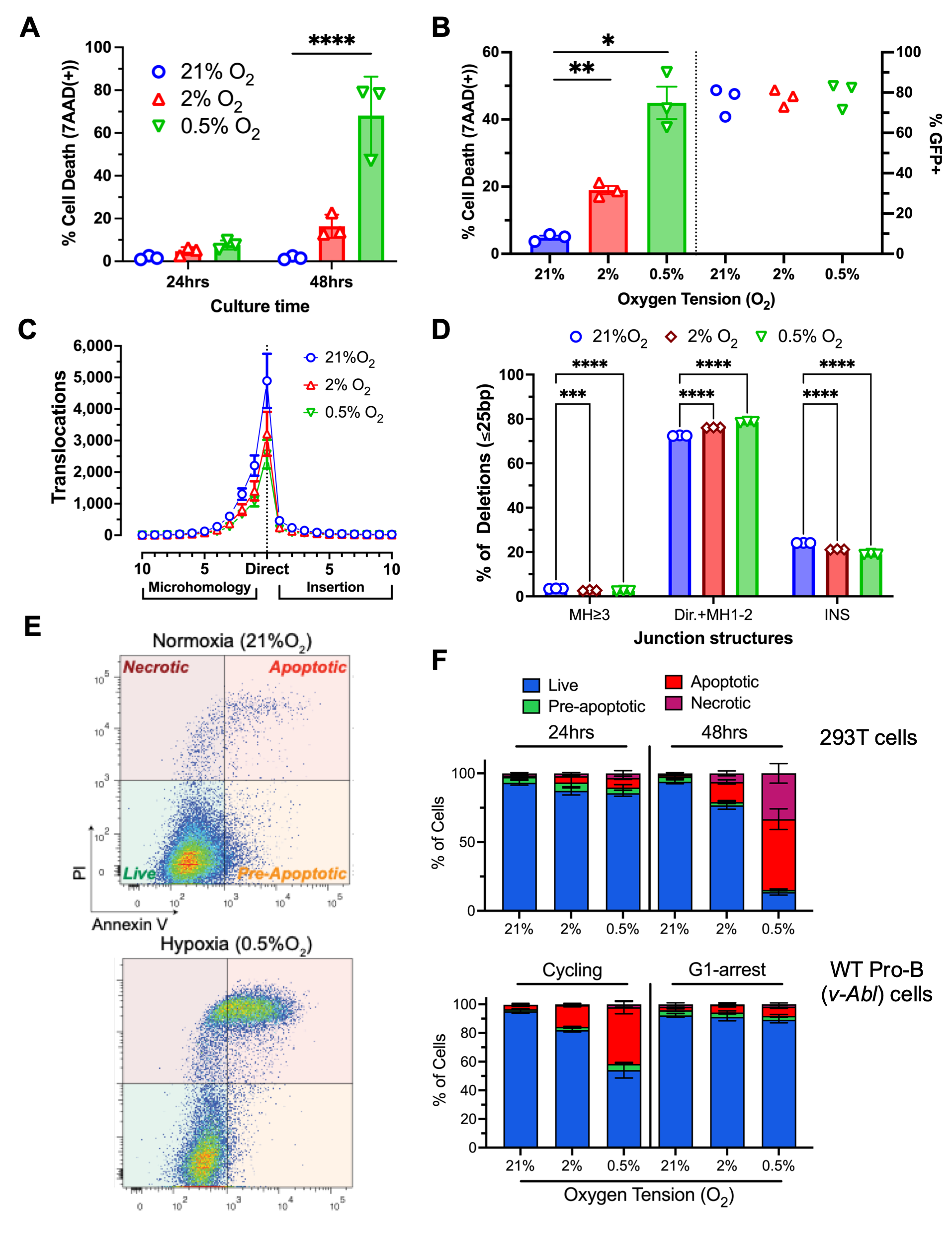

**Fig. S2: Hypoxia effects on viability and junction structures.** (A) Cell viability measures under chronic O_2_ tensions. (B) Cell viability and transfection of 293T cells for *JoinT-seq*. (C) Junction structures of translocations at varied O_2_ tensions. (D) Distributions of joint structures for small deletions; see Fig S1 legend for more details. Two-way ANOVA with Tukey posttest: ***P < 0.001, ****P < 0.0001. (E) Representative FACS plots measuring apoptosis via Annexin V. (F) Apoptosis measures for cycling 293T cells & both cycling and G1/G0-arrest progenitor B cell (*v-Abl*) lines.

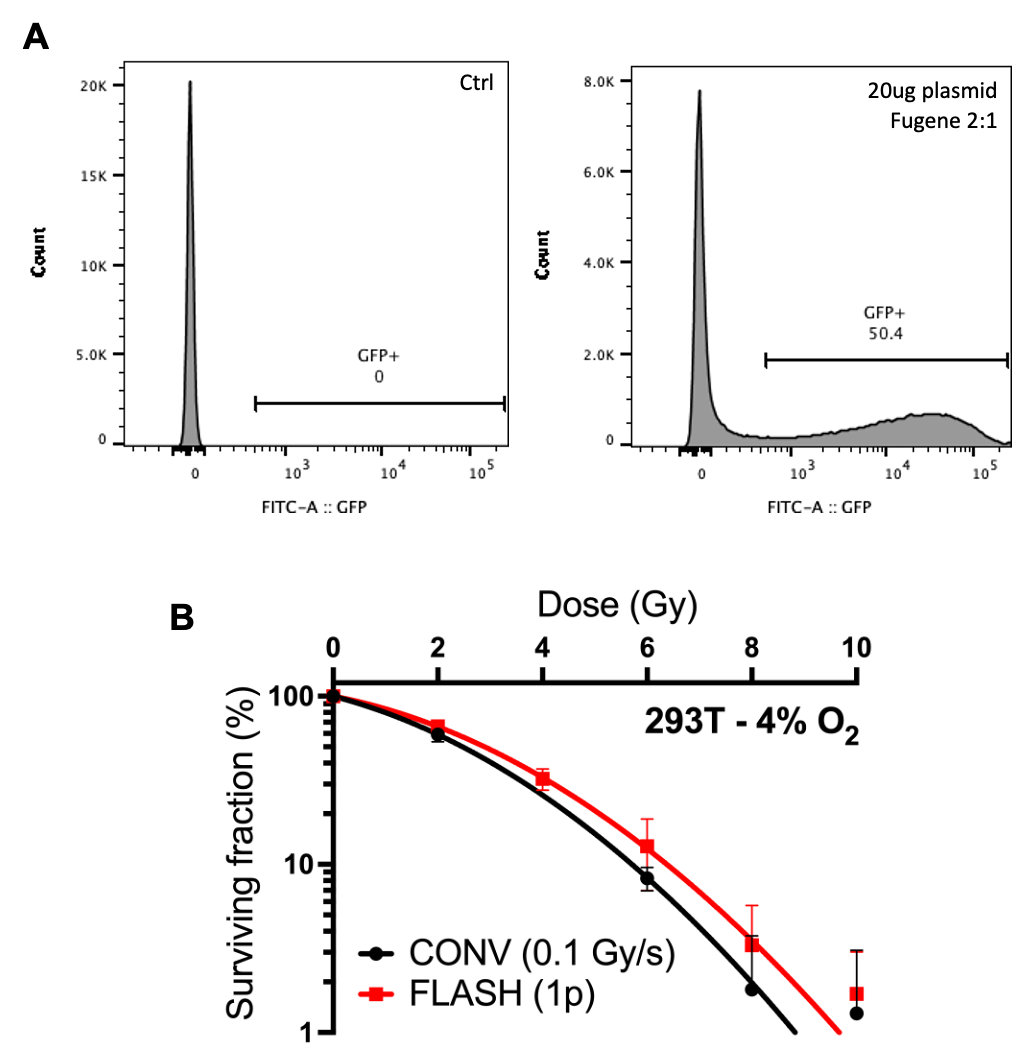

**Fig. S3: SpCas9:RAG1B plasmid transfection optimization and clonogenic survival after irradiation.** (A) GFP plasmid transfection using FuGENE (Promega). (B) Clonogenic survival analysis of CONV-RT or FLASH-RT 293T cells at 4% oxygen tensions.

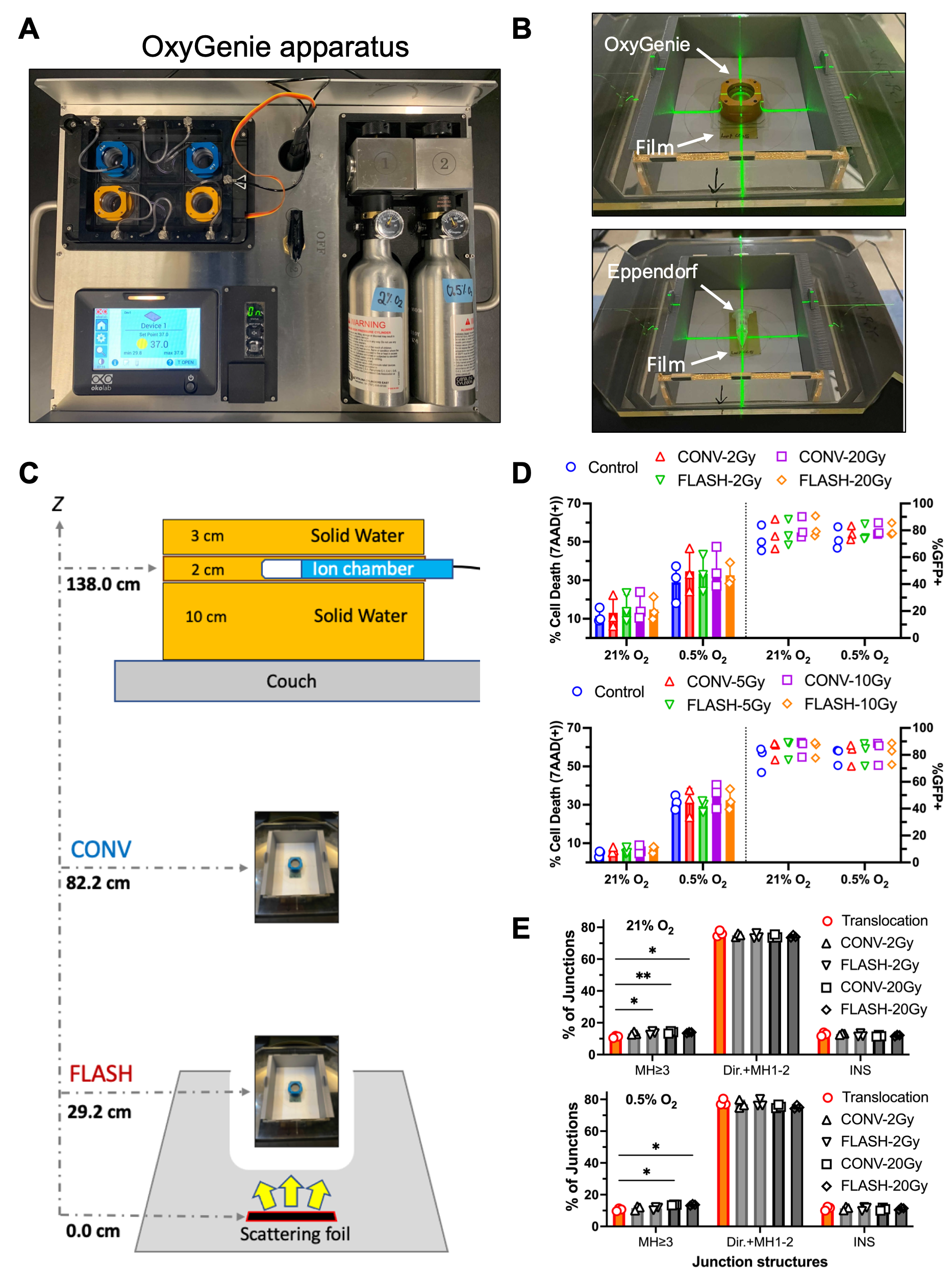

**Fig. S4: OxyGenie and CONV-RT vs FLASH-RT setup.** (A) Image of OxyGenie transportation apparatus. (B) Placement of samples (OxyGenie cassette or tube) in the Stanford LINAC. (C) LINAC configuration for CONV-RT and FLASH-RT delivery. (D) Cell viability and transfection efficiencies for biological replications (N=3). (E) Translocation junction structures; see Fig S1 legend for details. Two-way ANOVA with Tukey posttest: *P < 0.05, **P < 0.01.

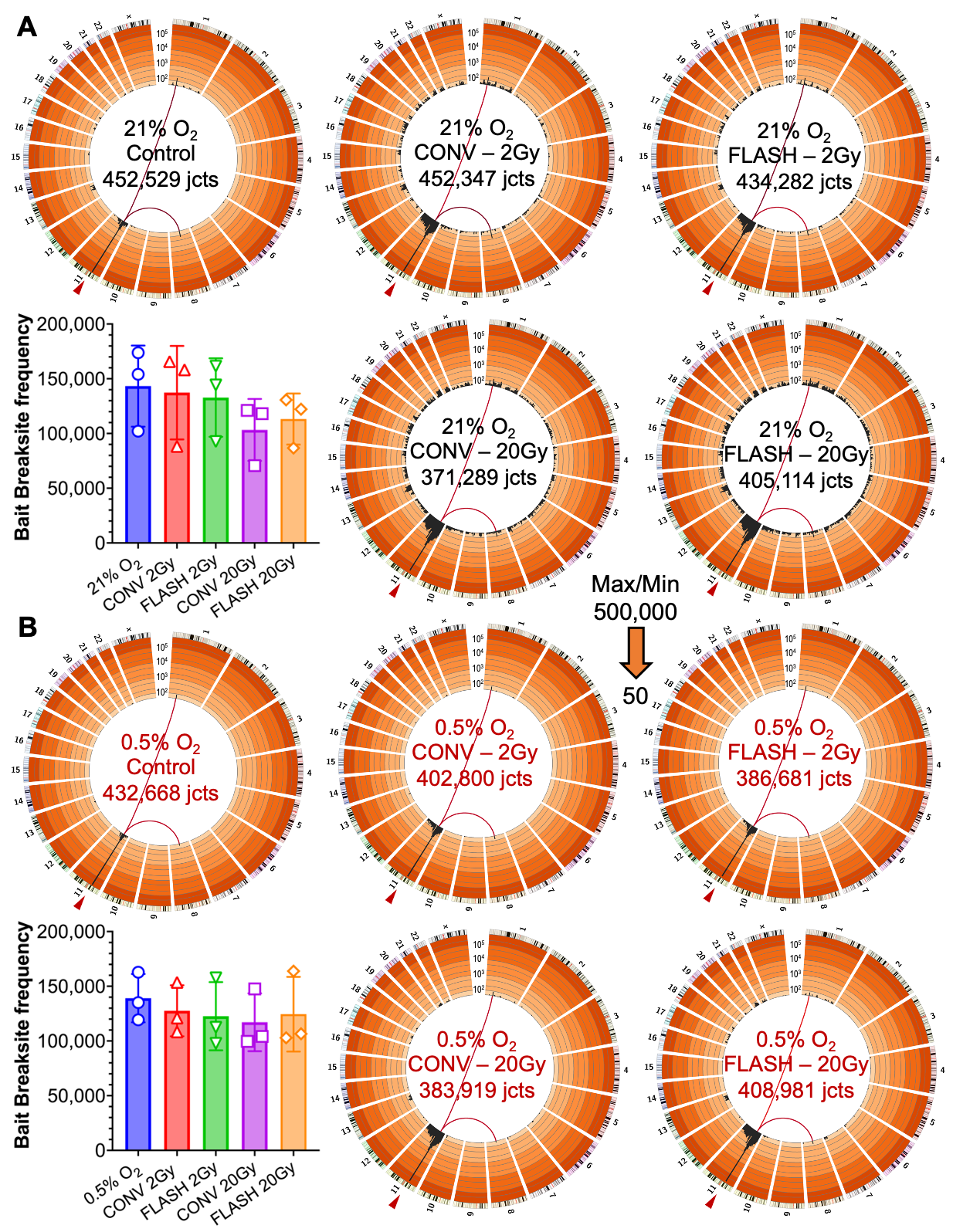

**Fig. S5: Circos plots and bait Breaksite frequencies for the 2 Gy / 20 Gy experiment.** (A) Circos plots and bait breaksite frequencies comparing CONV-RT vs. FLASH-RT (A) in air and (B) in hypoxia (N=3 each).

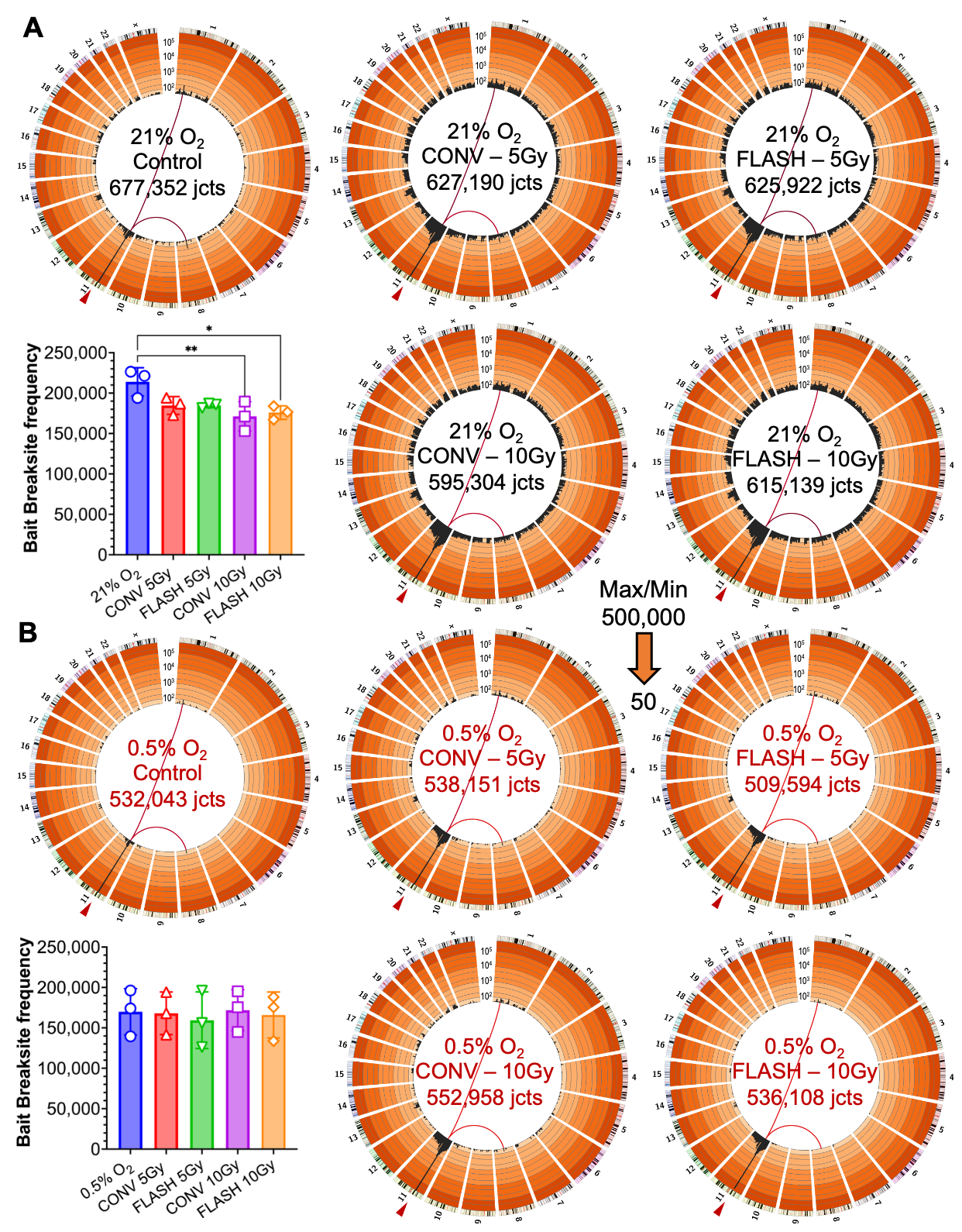

**Fig. S6: Circos plots and bait Breaksite frequencies for the 5 Gy / 10 Gy experiment.** (A) Circos plots and bait breaksite frequencies comparing CONV-RT vs. FLASH-RT (A) in air and (B) in hypoxia (N=3 each). One-way ANOVA with Dunnett’s post test: *P < 0.05, **P < 0.01.

**Table S1. *HTGTS-JoinT-seq* libraries information**

| **Irradiation** | | **Cas9 Bait**  ***** | | **O_2_%** | | | **Biol.**  **Rep** | | **Seq.**  **Reads** | | | | **Total Junctions** | | | **Breaksite junctions **** | | | **Intrachrom.**  **Transloc.**** | | | **Interchrom. Transloc.** | | | **% combined**  **Transloc.** |
| --- | --- | --- | --- | --- | --- | --- | --- | --- | --- | --- | --- | --- | --- | --- | --- | --- | --- | --- | --- | --- | --- | --- | --- | --- | --- |
| **Figure 1** | | | | |  | | |  | | |  | | |  | | |  | | |  | | |  | | |
| Control  (no IR) | | SaCas9:RAG1.1 | | 21% | | | 1 | | 904,846 | | | | 173,466 | | | 165,147 | | | 729 | | | 7,590 | | | 4.80% |
|  |  | SaCas9:RAG1.1 | | 21% | | | 2 | | 904,846 | | | | 180,450 | | | 170,832 | | | 802 | | | 8,816 | | | 5.33% |
|  |  | SaCas9:RAG1.1 | | 21% | | | 3 | | 904,846 | | | | 174,605 | | | 162,824 | | | 884 | | | 10,897 | | | 6.75% |
| CONV  10 Gy | | SaCas9:RAG1.1 | | 21% | | | 1 | | 904,846 | | | | 145,410 | | | 125,112 | | | 6,758 | | | 13,540 | | | 13.96% |
|  |  | SaCas9:RAG1.1 | | 21% | | | 2 | | 904,846 | | | | 145,154 | | | 123,577 | | | 6,997 | | | 14,580 | | | 14.86% |
|  |  | SaCas9:RAG1.1 | | 21% | | | 3 | | 904,846 | | | | 175,728 | | | 150,451 | | | 6,649 | | | 18,628 | | | 14.38% |
| **Figure 2** | | | | |  | | |  | | |  | | |  | | |  | | |  | | |  | | |
| Control  (No IR) | | SaCas9:RAG1.1 | | 21% | | | 1 | | 904,846 | | | | 206,619 | | | 194,360 | | | 893 | | | 11,366 | | | 5.93% |
|  |  | SaCas9:RAG1.1 | | 21% | | | 2 | | 904,846 | | | | 186,443 | | | 178,693 | | | 694 | | | 7,056 | | | 4.16% |
|  |  | SaCas9:RAG1.1 | | 21% | | | 3 | | 904,846 | | | | 200,789 | | | 188,034 | | | 907 | | | 11,848 | | | 6.35% |
| No IR | | SaCas9:RAG1.1 | | 2% | | | 1 | | 904,846 | | | | 183,442 | | | 174,929 | | | 591 | | | 7,922 | | | 4.64% |
|  |  | SaCas9:RAG1.1 | | 2% | | | 2 | | 904,846 | | | | 175,559 | | | 169,890 | | | 457 | | | 5,212 | | | 3.23% |
|  |  | SaCas9:RAG1.1 | | 2% | | | 3 | | 904,846 | | | | 160,786 | | | 154,143 | | | 427 | | | 6,216 | | | 4.13% |
| No IR | | SaCas9:RAG1.1 | | 0.5% | | | 1 | | 904,846 | | | | 169,155 | | | 162,714 | | | 459 | | | 5,982 | | | 3.81% |
|  |  | SaCas9:RAG1.1 | | 0.5% | | | 2 | | 904,846 | | | | 166,336 | | | 161,520 | | | 463 | | | 4,353 | | | 2.90% |
|  |  | SaCas9:RAG1.1 | | 0.5% | | | 3 | | 904,846 | | | | 112,818 | | | 107,803 | | | 320 | | | 4,695 | | | 4.45% |
| **Figure 3** | | | | |  | | |  | | |  | | |  | | |  | | |  | | |  | | |
| Control  (no IR) | | SpCas9:RAG1B | | 4% | | | 1 | | 214,700 | | | | 26,104 | | | 24,733 | | | 215 | | | 1,156 | | | 5.25% |
|  |  | SpCas9:RAG1B | | 4% | | | 2 | | 214,700 | | | | 22,419 | | | 21,445 | | | 179 | | | 795 | | | 4.34% |
|  |  | SpCas9:RAG1B | | 4% | | | 3 | | 214,700 | | | | 21,635 | | | 20,272 | | | 143 | | | 1,220 | | | 6.30% |
| X-Ray  10Gy | | SpCas9:RAG1B | | 4% | | | 1 | | 214,700 | | | | 15,391 | | | 8,902 | | | 2,609 | | | 3,880 | | | 42.16% |
|  |  | SpCas9:RAG1B | | 4% | | | 2 | | 214,700 | | | | 14,853 | | | 10,019 | | | 1,891 | | | 2,943 | | | 32.55% |
|  |  | SpCas9:RAG1B | | 4% | | | 3 | | 214,700 | | | | 22,578 | | | 16,772 | | | 2,104 | | | 3,702 | | | 25.72% |
| CONV  10 Gy | | SpCas9:RAG1B | | 4% | | | 1 | | 214,700 | | | | 18,250 | | | 12,579 | | | 2,025 | | | 3,646 | | | 31.07% |
|  |  | SpCas9:RAG1B | | 4% | | | 2 | | 214,700 | | | | 12,930 | | | 9,598 | | | 1,282 | | | 2,050 | | | 25.77% |
|  |  | SpCas9:RAG1B | | 4% | | | 3 | | 214,700 | | | | 21,791 | | | 16,835 | | | 1,732 | | | 3,224 | | | 22.74% |
| FLASH  10 Gy | | SpCas9:RAG1B | | 4% | | | 1 | | 214,700 | | | | 15,392 | | | 10,589 | | | 1,662 | | | 3,141 | | | 31.20% |
|  |  | SpCas9:RAG1B | | 4% | | | 2 | | 214,700 | | | | 13,458 | | | 9,004 | | | 1,698 | | | 2,756 | | | 33.10% |
|  |  | SpCas9:RAG1B | | 4% | | | 3 | | 214,700 | | | | 19,423 | | | 15,130 | | | 1,372 | | | 2,921 | | | 22.10% |
| **Figure 4, S4, S5** | | | | |  | | |  | | |  | | |  | | |  | | |  | | |  | | |
| Control  (no IR) | | SaCas9:RAG1.1 | | 21% | | | 1 | | 904,846 | | | | 106,901 | | | 102,121 | | | 607 | | | 4,173 | | | 4.47% |
|  |  | SaCas9:RAG1.1 | | 21% | | | 2 | | 904,846 | | | | 182,181 | | | 173,831 | | | 930 | | | 7,420 | | | 4.58% |
|  |  | SaCas9:RAG1.1 | | 21% | | | 3 | | 904,846 | | | | 163,447 | | | 154,084 | | | 795 | | | 8,568 | | | 5.73% |
| CONV  2 Gy | | SaCas9:RAG1.1 | | 21% | | | 1 | | 904,846 | | | | 96,783 | | | 88,188 | | | 2,008 | | | 6,587 | | | 8.88% |
|  |  | SaCas9:RAG1.1 | | 21% | | | 2 | | 904,846 | | | | 181,579 | | | 165,656 | | | 3,698 | | | 12,225 | | | 8.77% |
|  |  | SaCas9:RAG1.1 | | 21% | | | 3 | | 904,846 | | | | 173,985 | | | 158,154 | | | 3,112 | | | 12,719 | | | 9.10% |
| FLASH  2 Gy | | SaCas9:RAG1.1 | | 21% | | | 1 | | 904,846 | | | | 100,979 | | | 92,694 | | | 1,882 | | | 6,403 | | | 8.20% |
|  |  | SaCas9:RAG1.1 | | 21% | | | 2 | | 904,846 | | | | 175,437 | | | 161,593 | | | 3,147 | | | 10,697 | | | 7.89% |
|  |  | SaCas9:RAG1.1 | | 21% | | | 3 | | 904,846 | | | | 157,866 | | | 144,351 | | | 2,435 | | | 11,080 | | | 8.56% |
| CONV  20 Gy | | SaCas9:RAG1.1 | | 21% | | | 1 | | 904,846 | | | | 84,264 | | | 70,659 | | | 5,566 | | | 8,039 | | | 16.15% |
|  |  | SaCas9:RAG1.1 | | 21% | | | 2 | | 904,846 | | | | 143,908 | | | 118,126 | | | 10,477 | | | 15,305 | | | 17.92% |
|  |  | SaCas9:RAG1.1 | | 21% | | | 3 | | 904,846 | | | | 143,117 | | | 121,075 | | | 7,947 | | | 14,095 | | | 15.40% |
| FLASH  20 Gy | | SaCas9:RAG1.1 | | 21% | | | 1 | | 904,846 | | | | 104,748 | | | 86,741 | | | 7,238 | | | 10,769 | | | 17.19% |
|  |  | SaCas9:RAG1.1 | | 21% | | | 2 | | 904,846 | | | | 155,125 | | | 130,670 | | | 9,955 | | | 14,500 | | | 15.76% |
|  |  | SaCas9:RAG1.1 | | 21% | | | 3 | | 904,846 | | | | 145,241 | | | 122,164 | | | 8,297 | | | 14,780 | | | 15.89% |
| Control  (no IR) | | SaCas9:RAG1.1 | | 0.5% | | | 1 | | 904,846 | | | | 122,833 | | | 119,354 | | | 479 | | | 3,000 | | | 2.83% |
|  |  | SaCas9:RAG1.1 | | 0.5% | | | 2 | | 904,846 | | | | 167,997 | | | 162,886 | | | 595 | | | 4,516 | | | 3.04% |
|  |  | SaCas9:RAG1.1 | | 0.5% | | | 3 | | 904,846 | | | | 141,838 | | | 135,133 | | | 645 | | | 6,060 | | | 4.73% |
| CONV  2 Gy | | SaCas9:RAG1.1 | | 0.5% | | | 1 | | 904,846 | | | | 113,134 | | | 108,351 | | | 1,046 | | | 3,737 | | | 4.23% |
|  |  | SaCas9:RAG1.1 | | 0.5% | | | 2 | | 904,846 | | | | 160,313 | | | 153,509 | | | 1,279 | | | 5,525 | | | 4.24% |
|  |  | SaCas9:RAG1.1 | | 0.5% | | | 3 | | 904,846 | | | | 129,353 | | | 121,022 | | | 1,453 | | | 6,878 | | | 6.44% |
| FLASH  2 Gy | | SaCas9:RAG1.1 | | 0.5% | | | 1 | | 904,846 | | | | 117,540 | | | 112,539 | | | 1,068 | | | 3,933 | | | 4.25% |
|  |  | SaCas9:RAG1.1 | | 0.5% | | | 2 | | 904,846 | | | | 164,399 | | | 157,668 | | | 1,283 | | | 5,448 | | | 4.09% |
|  |  | SaCas9:RAG1.1 | | 0.5% | | | 3 | | 904,846 | | | | 104,742 | | | 97,865 | | | 1,093 | | | 5,784 | | | 6.57% |
| CONV  20 Gy | | SaCas9:RAG1.1 | | 0.5% | | | 1 | | 904,846 | | | | 108,777 | | | 99,705 | | | 3,115 | | | 5,957 | | | 8.34% |
|  |  | SaCas9:RAG1.1 | | 0.5% | | | 2 | | 904,846 | | | | 160,192 | | | 147,596 | | | 4,256 | | | 8,340 | | | 7.86% |
|  |  | SaCas9:RAG1.1 | | 0.5% | | | 3 | | 904,846 | | | | 114,950 | | | 104,134 | | | 3,447 | | | 7,369 | | | 9.41% |
| FLASH  20 Gy | | SaCas9:RAG1.1 | | 0.5% | | | 1 | | 904,846 | | | | 112,507 | | | 102,933 | | | 3,307 | | | 6,267 | | | 8.51% |
|  |  | SaCas9:RAG1.1 | | 0.5% | | | 2 | | 904,846 | | | | 177,265 | | | 163,866 | | | 4,406 | | | 8,993 | | | 7.56% |
|  |  | SaCas9:RAG1.1 | | 0.5% | | | 3 | | 904,846 | | | | 119,209 | | | 106,623 | | | 4,281 | | | 8,306 | | | 10.56% |
| **Figure 4, S4, S6** | | | | |  | | |  | | |  | | |  | | |  | | |  | | |  | | |
| Control  (no IR) | | SaCas9:RAG1.1 | | 21% | | | 1 | | 904,846 | | | | 230,570 | | | 221,371 | | | 860 | | | 8,339 | | | 3.99% |
|  |  | SaCas9:RAG1.1 | | 21% | | | 2 | | 904,846 | | | | 238,851 | | | 226,437 | | | 1,035 | | | 11,379 | | | 5.20% |
|  |  | SaCas9:RAG1.1 | | 21% | | | 3 | | 904,846 | | | | 207,931 | | | 194,002 | | | 1,082 | | | 12,847 | | | 6.70% |
| CONV  Gy | | SaCas9:RAG1.1 | | 21% | | | 1 | | 904,846 | | | | 208,782 | | | 187,002 | | | 6,002 | | | 15,778 | | | 10.43% |
|  |  | SaCas9:RAG1.1 | | 21% | | | 2 | | 904,846 | | | | 224,108 | | | 194,268 | | | 8,112 | | | 21,728 | | | 13.32% |
|  |  | SaCas9:RAG1.1 | | 21% | | | 3 | | 904,846 | | | | 194,300 | | | 172,788 | | | 4,575 | | | 16,937 | | | 11.07% |
| FLASH  5 Gy | | SaCas9:RAG1.1 | | 21% | | | 1 | | 904,846 | | | | 208,335 | | | 185,805 | | | 6,269 | | | 16,261 | | | 10.81% |
|  |  | SaCas9:RAG1.1 | | 21% | | | 2 | | 904,846 | | | | 209,317 | | | 181,542 | | | 7,548 | | | 20,227 | | | 13.27% |
|  |  | SaCas9:RAG1.1 | | 21% | | | 3 | | 904,846 | | | | 208,270 | | | 187,068 | | | 4,594 | | | 16,608 | | | 10.18% |
| CONV  10 Gy | | SaCas9:RAG1.1 | | 21% | | | 1 | | 904,846 | | | | 177,452 | | | 153,125 | | | 8,117 | | | 16,210 | | | 13.71% |
|  |  | SaCas9:RAG1.1 | | 21% | | | 2 | | 904,846 | | | | 204,017 | | | 171,094 | | | 10,705 | | | 22,218 | | | 16.14% |
|  |  | SaCas9:RAG1.1 | | 21% | | | 3 | | 904,846 | | | | 213,835 | | | 189,490 | | | 6,432 | | | 17,913 | | | 11.38% |
| FLASH  10 Gy | | SaCas9:RAG1.1 | | 21% | | | 1 | | 904,846 | | | | 195,578 | | | 167,243 | | | 9,544 | | | 18,791 | | | 14.49% |
|  |  | SaCas9:RAG1.1 | | 21% | | | 2 | | 904,846 | | | | 210,604 | | | 176,549 | | | 10,781 | | | 23,274 | | | 16.17% |
|  |  | SaCas9:RAG1.1 | | 21% | | | 3 | | 904,846 | | | | 208,957 | | | 183,882 | | | 6,490 | | | 18,585 | | | 12.00% |
| Control  (no IR) | | SaCas9:RAG1.1 | | 0.5% | | | 1 | | 904,846 | | | | 203,561 | | | 196,255 | | | 710 | | | 6,596 | | | 3.59% |
|  |  | SaCas9:RAG1.1 | | 0.5% | | | 2 | | 904,846 | | | | 181,326 | | | 174,066 | | | 658 | | | 6,602 | | | 4.00% |
|  |  | SaCas9:RAG1.1 | | 0.5% | | | 3 | | 904,846 | | | | 147,156 | | | 139,501 | | | 564 | | | 7,091 | | | 5.20% |
| CONV  5 Gy | | SaCas9:RAG1.1 | | 0.5% | | | 1 | | 904,846 | | | | 205,675 | | | 194,271 | | | 2,389 | | | 9,015 | | | 5.54% |
|  |  | SaCas9:RAG1.1 | | 0.5% | | | 2 | | 904,846 | | | | 181,251 | | | 168,142 | | | 2,574 | | | 10,535 | | | 7.23% |
|  |  | SaCas9:RAG1.1 | | 0.5% | | | 3 | | 904,846 | | | | 151,225 | | | 141,635 | | | 1,651 | | | 7,939 | | | 6.34% |
| FLASH  5 Gy | | SaCas9:RAG1.1 | | 0.5% | | | 1 | | 904,846 | | | | 207,627 | | | 196,020 | | | 2,468 | | | 9,139 | | | 5.59% |
|  |  | SaCas9:RAG1.1 | | 0.5% | | | 2 | | 904,846 | | | | 167,308 | | | 156,114 | | | 2,538 | | | 8,656 | | | 6.69% |
|  |  | SaCas9:RAG1.1 | | 0.5% | | | 3 | | 904,846 | | | | 134,659 | | | 126,191 | | | 1,500 | | | 6,968 | | | 6.29% |
| CONV  10 Gy | | SaCas9:RAG1.1 | | 0.5% | | | 1 | | 904,846 | | | | 188,077 | | | 175,488 | | | 3,092 | | | 9,497 | | | 6.69% |
|  |  | SaCas9:RAG1.1 | | 0.5% | | | 2 | | 904,846 | | | | 209,801 | | | 195,187 | | | 3,888 | | | 10,726 | | | 6.97% |
|  |  | SaCas9:RAG1.1 | | 0.5% | | | 3 | | 904,846 | | | | 155,080 | | | 144,905 | | | 2,159 | | | 8,016 | | | 6.56% |
| FLASH  10 Gy | | SaCas9:RAG1.1 | | 0.5% | | | 1 | | 904,846 | | | | 202,700 | | | 188,255 | | | 3,705 | | | 10,740 | | | 7.13% |
|  |  | SaCas9:RAG1.1 | | 0.5% | | | 2 | | 904,846 | | | | 189,548 | | | 175,572 | | | 3,777 | | | 10,199 | | | 7.37% |
|  |  | SaCas9:RAG1.1 | | 0.5% | | | 3 | | 904,846 | | | | 143,860 | | | 133,815 | | | 2,245 | | | 7,800 | | | 6.98% |

* SaCas9:RAG1.1 bait DSB – chr11: 36,573,331; SpCas9:RAG1B bait DSB – chr11: 36,573,286

** Translocations are defined as junctions outside of the 500kb flanking bait breaksite region

**Table S2. Electron Beam irradiation parameters for *HTGTS-JoinT-seq***

|  | Varian Trilogy LINAC  (Stanford) | | | | | | Oriatron eRT6 LINAC  (CHUV) | | |
| --- | --- | --- | --- | --- | --- | --- | --- | --- | --- |
| Modality | | CONV | FLASH | | | CONV | | FLASH | |
| Beam energy [MeV] | 15.7 | | | 16.6 | 5.5 | | | | 5.5 |
| Target dose [Gy] | 2, 5, 10, 20 | | | 2, 5, 10, 20 | 10 | | | | 10 |
| Delivered pulses | 1901, 4763, 9526, 19051 | | | 2, 5, 10, 20 | 477-482 | | | | 2 |
| Dose per pulse [Gy] | 1.05E-3 | | | 1 | 0.02 | | | | 5 |
| Pulse rate [pulses/sec] | 72 | | | 90 | 10 | | | | 100 |
| Dose rate [Gy/sec] | 0.08 | | | 180, 112.5, 100, 94.8 | 0.21 | | | | 1,000 |
| Pulse length/width [μsec] | 3.75 | | | 3.75 | 1.0 | | | | 2.1 |
| Intra-pulse dose rate [Gy/μsec] | 2.80E-4 | | | 0.27 | 0.02 | | | | 2.38 |
| Delivery time [sec] | 26.4, 66.2, 132.2, 264.6 | | | 0.01, 0.04, 0.10, 0.21 | 48.20 | | | | 0.01 |

**Table S3. Electron Beam irradiation parameters for clonogenic survival**

|  | **Oriatron eRT6 LINAC**  **(CHUV)** | | | |
| --- | --- | --- | --- | --- |
| **Modality** | | CONV | FLASH | |
| **Beam energy (MeV)** | 5.5 | | | 5.5 |
| **Target dose [Gy]** | 2, 4, 6, 8 | | | 2, 4, 6, 8 |
| **Delivered pulses** | 136, 272, 408, 540 | | | 1 |
| **Dose per pulse [Gy]** | 0.02 | | | 2, 4, 6, 8 |
| **Pulse rate [pulses/sec]** | 10 | | | 100 |
| **Dose rate [Gy/sec]** | 0.15 | | | 1.1E6 – 4.4E6 |
| **Pulse length/width [μsec]** | 1.0 | | | 1.8 |
| **Intra-pulse dose rate [Gy/μsec]** | 0.02 | | | 1.1E6 – 4.4E6 |
| **Delivery time [sec]** | 13.5, 27.1, 40.7, 53.9 | | | 1.80E-6 |
